# Control of invasive *Phragmites australis* (European common reed) alters macroinvertebrate communities

**DOI:** 10.1101/2021.08.25.457681

**Authors:** C.D. Robichaud, J.V. Basso, R.C. Rooney

## Abstract

Wetland restoration often involves invasive plant species suppression to encourage the recovery of native-dominated vegetation communities. However, assessment of recovery is usually focused only on vegetation and the response of other critical wetland biota, such as macroinvertebrates, is seldom assessed. We characterized the aquatic, semi-aquatic, and terrestrial macroinvertebrate communities in remnant, uninvaded marsh to identify restoration targets and compared this to the communities in *Phragmites australis-invaded* marsh, and in formerly invaded marsh that was treated with the herbicide glyphosate in 2016 to simultaneously evaluate the effects of invasion and of invasive species suppression. We sampled invertebrates in 2017 and 2018 to track two years following herbicide treatment. The invertebrate community composition captured by the emergence traps was similar between *P. australis* and remnant marsh, suggesting invasion has little effect on macroinvertebrate community structure. There was also high concordance between the aquatic and emerging invertebrate communities in the invaded and uninvaded habitats. In contrast, herbicide-treated sites had a unique community composition, characterized by very high densities of Chironomidae (Diptera) and low taxa richness and evenness. Herbicide-treated sites also exhibited low concordance between the aquatic and emerging invertebrate communities, potentially attributable to the sparse emerging vegetation cover providing limited substrates for emergence. Herbicide-based invasive species control results in considerable changes to the macroinvertebrate community in freshwater marshes for at least two years after treatment, which may have consequences for aquatic food webs and species that rely on macroinvertebrates as prey.

## Introduction

Invasive plant species pose a global threat to biological communities as they can alter biogeochemical cycling regimes (Vilà et al. 2011), hydrology (Le Maitre 2004), and habitat structure (Crooks 2002) which negatively effects native species. Often, invasive species are associated with degraded environments (e.g., Fruh et al. 2012) and benefit from stressors, such as pollution and altered disturbance regimes, while also driving declines in native species and further ecological impairment (e.g., MacDougall & Turkington 2005; Bauer 2012). Ecological restoration of degraded environments is frequently hampered by the presence of established invasive species (D’Antonio & Meyerson 2002). Typically, the first step in restoring these degraded ecosystems is the control or removal of an invasive plant, with the intention that the native biological community will reassemble (Weidlich et al. 2020). However, often studies following the removal of an invasive plant species only measure variables related to the vegetation community recovery (e.g., Hazelton et al. 2014). To fully assess the effects of invasive species removal and restoration efforts it is essential to compare the biological communities present in restored habitats to communities in both 1) invaded vegetation and 2) remnant, uninvaded vegetation. Otherwise, restoration targets are undefined and restoration outcomes cannot be evaluated.

Wetlands are particularly vulnerable to invasive plants due to their dynamic environmental conditions (Zedler & Kercher 2004) and ongoing global loss and degradation (e.g., Brinson & Malvárez 2021). In North America, Lake Erie has approximately 19,330 ha of coastal wetlands which are some of the most diverse and species rich in the Laurentian Great Lakes region (Herdendorf 1992; Ball et al. 2003). This diversity includes complex macroinvertebrate communities consisting of aquatic, semi-aquatic, and terrestrial taxa (Batzer et al. 1999). Macroinvertebrates are often the most diverse and abundant fauna within a wetland and provide an essential link between primary producers (e.g., plants, algae) and higher trophic levels (Batzer et al. 1999). Wetland macroinvertebrates are resilient to natural variations in their environment but are likely to respond strongly to dramatic changes (Batzer 2013, Gathman and Burton 2011). In Lake Erie coastal wetlands, this poses an interesting hypothesis as wetlands are changing dramatically due to invasive plant species and ongoing management to suppress invasive plant species.

Invasive *Phragmites australis* subsp. *australis* (European common reed [(Cav.) Trin. Ex Steud.]) is a perennial wetland grass that has established throughout Great Lake wetlands (e.g., Wilcox 2012). *Phragmites australis* rapidly expanded in Lake Erie beginning in the late-1990s after periods of low water levels (Tulbure & Johnston 2010; Wilcox 2012). Within three decades, *P. australis* had created monocultures in formerly diverse wetland systems including the Long Point peninsula, a World Biosphere Reserve on Lake Erie (Wilcox et al. 2003; Jung et al. 2017). Monocultures of *P. australis* produce tall, dense stands and large amounts of litter (e.g., Yuckin & Rooney 2019), decrease wetland avian diversity (e.g., Robichaud & Rooney 2017), and are detrimental to endangered reptiles (e.g., Markle & Chow-Fraser 2018) and amphibians (e.g., Greenberg & Green 2013). The documented negative effects of invasive *P. australis* have led to on-going efforts to control populations and restore native wetland vegetation in North America (e.g., Hazelton et al. 2014). This includes the recent large-scale (>3400 ha) application of a glyphosate-based herbicide directly over standing water in the marshes of Long Point and Rondeau Bays, an unprecedented control action in Canada (Robichaud & Rooney 2021a). Herbicide-application is a common tool used to manage *P. australis* (e.g., Hazelton et al. 2014), and while glyphosate-based herbicide degrades rapidly to non-toxic levels (Robichaud & Rooney 2021b) and is not likely a toxicological concern for aquatic macroinvertebrates, large macrophyte die-offs can affect native fauna (e.g., Linz et al. 1999).

Published studies report conflicting responses of macroinvertebrate density and diversity to *P. australis* invasion and treatment in North America, likely due to the variety of invertebrate sampling techniques and environmental conditions of invaded wetlands. In freshwater marshes, benthic macroinvertebrate density has been noted as similar among *P. australis* and other vegetation (e.g., *Typha angustifolia* (narrow-leaved cattail) or native flora) (Holomuzki & Klarer 2010) or higher in *P. australis* and treated *P. australis* due to gastropod and Chironomidae abundance (Kulesza et al. 2008). In brackish marshes, macroinvertebrate density and richness were higher in *Spartina* (cord-grass) vegetation than *P. australis* when sampled with soil cores and litter packs (Angradi et al. 2001). Assessing the invertebrate community present in standing vegetation in brackish marshes determined the functional traits of the invertebrate community in *P. australis* were significantly different than those using native vegetation, but recovered after *P. autralis* populations were controlled (Gratton & Denno 2005). In salt marshes, comparisons of the density and diversity of macroinvertebrates in *P. australis* and other vegetation often provide conflicting results depending on the sampling approach (e.g., sediment, litter, or pitfall traps) (Talley & Levin 2001; Fell et al. 2006). Most recent, a study contrasting three types of native vegetation community with *P. australis-invaded* marsh around the Great Salt Lake in Utah concluded that only pickleweed-dominated native vegetation supported a distinctive arthropod community (Leonard et al. 2021). This lack of consistency in invertebrate response to *P. australis*-invasion and its removal thus transcends environment (freshwater, brackish or saltwater) and micro-habitat (benthic, epiphytic, litter).

Aquatic macroinvertebrates can also be useful indicators of ecosystem condition - they often have predictable responses to environmental stressors (Bonada et al. 2006) - though there are challenges selecting bioindicators for the wide natural range of variation in Great Lakes coastal wetlands (e.g., Batzer 2013). A critical subset of aquatic macroinvertebrates have a terrestrial life cycle stage, connecting aquatic and terrestrial ecosystems. Additionally, there are a diversity of semi-aquatic and terrestrial macroinvertebrates that rely on wetlands for habitat (Batzer & Wu 2020) and are important components of wetland food webs. Neglecting to sample emerging and terrestrial components of the wetland macroinvertebrate community might be partly responsible for the lack of consistency in results among published studies and may limit inferences regarding the effects of invasive plants and their control (Harvey et al. 2014). For example, direct application of herbicide to wetland vegetation has been reported to increase the biomass of emerging invertebrates, particularly detritivores, with consequences for aquatic-terrestrial linkages and food webs (e.g., Linz et al. 1999; Baker et al. 2014). Therefore, it is important to sample multiple microhabitats to fully capture changes resulting from invasion and invasive plant suppression.

We compared the macroinvertebrate community structure in *P. australis*-invaded marsh, recently herbicide-treated marsh, and remnant, uninvaded marsh to test for effects of *P. australis* invasion and removal on this key wetland food web component. We sampled macroinvertebrates using emergence traps and in the aquatic habitat quantitatively using water column and vegetation collection methods to capture aquatic, semi-aquatic, and terrestrial macroinvertebrates that rely on wetlands. We addressed three research questions: 1) Do *P. australis* stands have lower densities or taxonomic richness than remnant, uninvaded marsh? 2) Does the macroinvertebrate community in herbicide-treatment areas more closely resemble remnant uninvaded marsh, or *P. australis-invaded* marsh? and 3) Does the agreement between the juvenile aquatic macroinvertebrate community composition and the winged terrestrial community composition differ among *P. australis*-invaded marsh, remnant marsh, and herbicide-treated sites?

## Methods

### *Study Site and* P. australis *Removal*

Our study took place in Long Point, Ontario, Canada. Located on the north shore of Lake Erie, Long Point is a sandspit peninsula that contains ecologically significant vegetation communities and provides habitat for hundreds of species (Ball et al. 2003). Populations of invasive *Phragmites australis* subsp. *australis* have been growing in Long Point since the late 1990s, displacing resident wetland vegetation communities and replacing them with dense stands of *P. australis* (Wilcox et al. 2003). As there were documented negative effects of *P. australis* on species-at-risk in Long Point (e.g., Greenberg and Green 2013), land managers started wetland restoration efforts that began with controlling the large population of *P. australis.* This control involved using a glyphosate-based herbicide directly over standing water, a first in Canada, and required extensive permitting as there was no registered herbicide product at the time. The herbicide application was carried out under a Permit to Perform an Aquatic Extermination of invasive *P. australis* in standing water issued by the Ontario Ministry of Environment, Conservation and Parks and an Emergency Use Registration (no. 32356) issued by Health Canada’s Pest Management Regulation Authority under the Pest Control Products Act.

In 2016, the Ontario Ministry of Resources and Forestry and the Nature Conservancy of Canada treated over 300 ha of *P. australis* in Long Point by applying a glyphosate-based herbicide (Roundup^®^ Custom for Aquatic & Terrestrial Use Liquid Herbicide, Bayer CropScience, Whippany, New Jersey) via helicopter (Eurocopter A-Star, Marseille Provence Airport, Marignane, France) in late August. Secondary treatment to remove standing dead culms of *P. australis* took place over the winter, using a combination of rolling and cutting (full treatment details in Robichaud & Rooney, 2021a & b). Following the removal of *P. australis*, sites were left to passively recolonize. In companion papers, we describe the fate of this herbicide (Robichaud and Rooney 2021b) and the recovery of vegetation in these marshes in response to *P. australis* suppression activities (Robichaud & Rooney 2021a). As part of vegetation and glyphosate monitoring, we surveyed the vegetation and substrate properties of control and treatment regions of the study area both before herbicide was applied in 2016 and afterward through 2018. The key factors that structure macroinvertebrate communities, other than predators, include water depth, substrate properties, and vegetation structure (community composition, canopy height, open water, light penetration, stem density etc.; Gathman and Burton 2011, Batzer 2013). Because we found no difference in these factors between the control (untreated *P. australis*) sites and the treatment (herbicide-treated *P. australis*) sites in the study area before herbicide application in 2016 (Robichaud & Rooney 2021a), we expect that the macroinvertebrate communities were also similar among the herbicide-treated and *P. australis*-invaded sites in our macroinvertebrate study.

In 2017, the spring following herbicide application, we established 27 sampling locations in three habitat types (Fig. 1): *P. australis-invaded* habitat (n = 9), herbicide-treated habitat (n = 9) and remnant, uninvaded marsh vegetation (hereafter “remnant” marsh; n = 9). The different sites were spaced a minimum of 150 m apart to ensure they were independent from the perspective of invertebrates. Herbicide treatment in 2016 was restricted to the western portion of the study area, about 2 km west of the location of the other sampling sites but spanned the same gradient in water depth and substrate composition (Table S1). Prior to herbicide treatment in 2016, our baseline vegetation monitoring confirmed that the herbicide-treated and untreated *P. australis*-invaded sites covered the same range in stem density, canopy height, light penetration, and floristic diversity (Robichaud and Rooney 2021a) as well as the same sandy and organic substrate. Further, sites were selected to ensure the same range of water depths was incorporated for each of the three habitat types. We consequently do not consider the distance between the herbicide-treated sites and the other stations to confound any effects of herbicide application on the macroinvertebrate communities.

**Figure 1.**
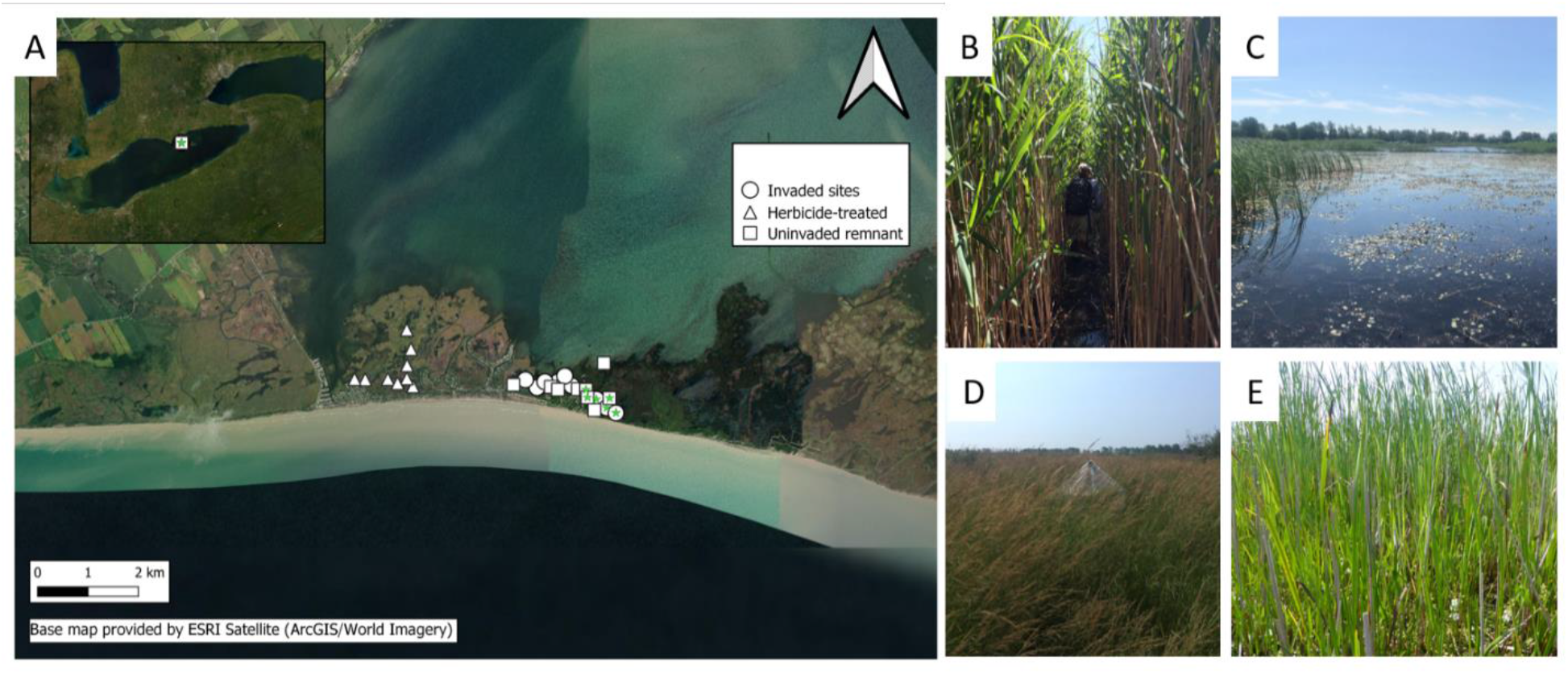
Emergence trap sites in Long Point, ON, Canada (A) situation in *P. australis* stands (B), herbicide-treated sites (C) and remnant marsh (D & E). Each site consisted of a modified emergence trap to capture aquatic, semi-aquatic, and terrestrial invertebrates. Sites were sampled once for aquatic invertebrates in May 2018. Emergence traps were sampled from June 19^th^ to July 21^st^ in 2017 and May 20^th^ to July 23^rd^ in 2018. New sites are marked with a star symbol and were established to compensate for sites that were herbicide-treated between 2017 and 2018 due to on-going *P. australis* control work in the park.

Although prior to herbicide application, the *P. australis-invaded* vegetation in the untreated and herbicide-treated sites were both heavily dominated by *P. australis* (mean vegetation richness = 4), with high stem density (ca. 100 total stems per m^2^), and tall canopies (mean = 3.8 m), following herbicide application, the vegetation community in treated sites was transformed. Herbicide-treated sites were characterized by open water with abundant submersed and floating aquatic vegetation (e.g., *Utricularia vulgaris, Potomogeton* spp.) and limited robust emergent vegetation (e.g., cattails, rushes) (Robichaud & Rooney 2021a). The gradient of water depth sampled within the uninvaded marsh captured an ecotone of remnant vegetation communities, from meadow marsh (characterized by sedges, grasses, and forbs, with shallow standing water (< 30 cm in May)) to emergent marsh (characterized by dense cattail (*Typha* spp.) and deeper standing water). Sites at intermediate water depth were characterized by a mix of meadow marsh and emergent marsh vegetation. Working alongside ongoing restoration projects provides realism for research, but sometimes at the expense of experimental design. Between 2017 and 2018, three of our *P. australis-invaded* sites were treated with herbicide as part of ongoing management and three resident vegetation (emergent marsh) sites were compromised by the passage of heavy machinery. We consequently replaced those six sites with new stations in the appropriate vegetation type for our 2018 surveys (Fig. 1).

### Field Methods

Beginning mid-June 2017, we deployed an emergence trap at each station (N = 27).

Emergence traps capture invertebrates with an aerial life stage to quantify the biomass of invertebrates per meter-squared. Each trap consisted of a capture vessel atop a pyramid structure on a 1 m^2^ base and covered in 500 μm mesh netting (following Anderson & Davis 2013; Figure S1). Traps ranged in height based on the vegetation community they were deployed in; 1.5 m tall in herbicide-treated and meadow marsh sites and 2 m tall in emergent marsh and *P. australis* invaded sites. When setting up the traps we made a concerted effort not to disrupt the surrounding vegetation so that we would not affect vegetation density or light penetration that could influence emergence timing or success. The capture vessel contained an aqueous solution of 70% ethanol as a preservative. It was collected and replaced approximately every 10 days from deployment until the end of July. This resulted in four collections in 2017 from 19-June-2017 to 21-July-2017. We repeated this in 2018, deploying traps in mid-May and collecting every 10 days on six occasions between 20-May-2018 to 23-July-2018. As described above, six stations had to be relocated between 2017 and 2018 due to disturbance of the original sites. Occasional storms damaged the traps or upset the collection vessels, resulting in missed collections from individual traps. The number of collections on each collection date is reported in Table S1.

At the end of July in 2017 and 2018, during peak aboveground plant biomass, we characterized the vegetation community composition and measured water depth, water temperature, and average canopy height at each site. We randomly placed three ¼ m^2^ quadrats within 10 m of the emergence traps and assessed the community composition of living and non-living cover (e.g., percent open water) using a modified Braun-Blanquet approach (e.g., Wikum & Shanholtzer 1978) such that each quadrat added up to 100% cover. We also measured water temperature (°C), water depth (cm), and canopy height (cm) within each quadrat. For each site, we averaged values from each quadrat to give one measure of community composition, water depth, water temperature, and canopy height.

Between 11 and 15 of May 2018 we sampled the aquatic macroinvertebrates to compare the community living in the water with the community sampled by our emergence traps. We used a ¼ m^2^ quadrat, within which we clipped and collected all standing dead litter, submersed, floating, and emergent vegetation into a bucket. We rinsed this vegetation with filtered lake water through a 500 μm mesh sieve, repeating this rinse process at least four times by adding filtered lake water, agitating the vegetation, and passing it through the sieve. The contents of the sieve were transferred to a collection jar and preserved in 70% ethanol. We then sorted through the remaining vegetation in white trays to collect any clinging invertebrates and added these to the collection jar. Note we also collected two replicate 10 cm diameter water column samples at each quadrat sampling location but did not include these in analyses because after processing 100% of the water column they yielded relatively low densities and no additional macroinvertebrate taxa. For clarity, we refer to emergence trap samples as “emergence trap invertebrates” and quadrat samples as “aquatic invertebrates” throughout.

### Laboratory Methods

Aquatic samples were sorted to separate macroinvertebrates from debris under a dissecting microscope. Individuals from both aquatic samples and emergence trap samples were identified to the lowest taxonomic level that did not require slide mounting, typically Order or Family, using Merritt et al. (2008), Thorp & Covich (1991), Triplehorn & Johnson (2005), and Marshall (2006). Descriptive statistics for wetland invertebrate communities generated using Family level ID are high congruent with the same statistics developed using Genus level (Piers et al 2021), thus we expect that our conclusions would be identical even with the additional effort of finer taxonomic resolution.

The two deepest sites in remnant and *P. australis*-invaded habitats are not represented in the aquatic invertebrate analyses as they were open water hemi-marsh and did not have adequate amounts of vegetation for sampling. Therefore, for emergence trap analyses we compared nine sites from each of the three habitat types, for aquatic invertebrate samples we compared nine herbicide-treated sites with eight sites in remnant marsh and eight sites in *P. australis*-invaded marsh. When comparing the emerging and aquatic invertebrate communities, we also excluded the emergence trap samples from the deepest remnant marsh and *P. australis*-invaded marsh, to maintain sampling balance. The process of sorting, identifying, and counting individuals took approximately two years (2018 – 2020) and yielded 28,744 individuals from aquatic samples and 47,772 individuals from emergence trap samples.

### Data Analysis

Taxa with two or fewer occurrences in the emergence trap samples across all sites and collection dates were removed from the emergence trap datasets and taxa with two or fewer occurrences in the aquatic invertebrate samples were removed from the aquatic invertebrate dataset to mitigate the influence of rare taxa on distance-based community analyses. The macroinvertebrate density from aquatic samples (1/4 m^2^ quadrats) were multiplied by four so the densities are comparable between the aquatic invertebrate samples and emergence trap samples (1 m^2^). The final taxa included in the aquatic invertebrate and emergence trap invertebrate analyses are reported in Table S2 and Table S3.

### Univariate analyses

As the macroinvertebrate data did not meet the assumptions of a parametric test, we performed permutational general linear models with taxonomic richness, density, and Pielou’s evenness (*J*) as response variables (calculated using the *vegan* package (Oksanen et al. 2020) in R v. 4.0.0 (R Core Team 2020)) and habitat type (*P. australis-invaded* marsh, remnant marsh, and herbicide-treated marsh) as a fixed factor for the aquatic invertebrate data, the 2017, and the 2018 emergence trap invertebrate data separately. Each test was run with 999 permutations, producing a p-value based on the number of test statistics from randomized runs that are as or more extreme than the actual test statistic. If habitat type was significant, we performed a Tukey’s post-hoc test. General linear modeling was performed using *lmPerm* (Wheeler & Torchiano 2016) and *agricolae* (de Mendiburu 2020) in R v. 4.0.0 (R Core Team 2020).

### Multivariate analyses

The following multivariate analyses were performed on the aquatic invertebrate, the 2017, and the 2018 emergence trap datasets separately. First, for the emergence trap data, we summed the densities of each taxon across all the visits in a given year (2017 = 4 visits, 2018 = 6 visits). For both the aquatic invertebrate data and the emergence trap data for each year, we relativized densities by the taxon’s annual maximum to reduce the influence of highly abundant taxa. Last, we calculated Bray-Curtis dissimilarity matrices (n = 3), which we used in subsequent multivariate analyses.

For each dissimilarity matrix, we used a permutation test (PERMDISP2) to assess differences in multivariate dispersion among the habitat types by measuring the mean distance to a group centroid in multivariate space (Anderson 2006; Anderson et al. 2006). To determine if the differences between groups were significant, we used a test that permuted the least-square residuals 999 times to generate a distribution to test the F-statistic against (Anderson et al. 2006). PERMDISP can provide information on beta diversity within each habitat type and determine if data meet the assumptions of homogeneity of variances necessary for a perMANOVA (Warton et al. 2012). As we were also interested in comparing the invertebrate community composition among the habitat types, we performed a perMANOVA (999 permutations) with habitat type as a fixed factor if homogeneity of variance assumptions were met.

We used a non-metric multidimensional scaling (NMDS) ordination to visualize differences in invertebrate community composition among the habitat types for each dissimilarity matrix. We included the site characteristics (vegetation community, water depth and temperature, and canopy height) as vectors in the three ordinations to help explain patterns in macroinvertebrate community composition. PERMDISP, perMANOVA, and NMDS ordinations were all performed using the *vegan* package (Oksanen et al. 2020) in R v. 4.0.0 (R Core Studios 2020). All figures were created with *ggplot* (Whickham 2016).

To model changes in the emergence trap invertebrate community over the sampling period in each year, we used multivariate generalized linear models. We ran two models, one for each year, with habitat type and “days since May 20” as fixed factors and a negative binomial distribution function. We then took the model output and calculated univariate test statistics and adjusted p-values for each Family to elucidate patterns for the different taxa. Analyses were performed using *mvabund* (Wang et al. 2021) in R v. 4.0.0.

### Congruence between aquatic and emerging invertebrates

We conducted congruence analyses on a reduced subset of macroinvertebrate taxa that had an aquatic larval stage and adult winged stage. Aquatic macroinvertebrate samples were taken in mid-May 2018 and compared to the macroinvertebrates collected from the emergence traps from 5 June to 23 July 2018 for congruence analyses. Because no aquatic macroinvertebrates were sampled from the deepest remnant and *P. australis*-invaded sites, we also excluded their emergence trap samples. This resulted in a total of 25 sites with paired aquatic and emerging macroinvertebrate communities. Because not all taxa captured in the aquatic invertebrate samples have a winged adult stage and not all macroinvertebrates captured by the emergence traps have an aquatic juvenile stage (i.e., some were terrestrial or semi-aquatic), we excluded taxa that did not have both an aquatic juvenile stage and a winged adult stage which resulted in 17 taxa in the aquatic invertebrate dataset and 31 in the emergence trap invertebrate dataset (Table S4). We then ran three Procrustes tests, one for each habitat type, using the protest function in *vegan* (Oksanen et al. 2020). This test performs a symmetric Procrustes analysis and permutations to estimate the significance of the t-statistic. We report the Procrustes correlation from the non-permuted solution and the p-value after 999 permutations. We also visualized the relationship between the aquatic and emerging macroinvertebrates among the three habitat types using an NMDS ordination.

## Results

### Univariate

#### Aquatic invertebrates

We detected a total of 29 taxa in *P. australis* stands, 26 taxa in herbicide-treated sites, and 37 taxa in remnant marsh (Table S5). Aquatic invertebrate density was significantly higher (permutational ANOVA F_2,22_ = 7.301, p = 0.004) in herbicide-treated sites than *P. australis-*invaded sites or remnant marsh habitat (Fig. 2A, Table 1). Two herbicide-treated sites had very high densities – 39,404 individuals per 1 m^2^ and another with 24,188 individuals per 1 m^2^ – consisting of high densities of Oligochaeta and Chironomidae. Pielou’s evenness was significantly lower in *P. australis-invaded* habitat (permutational ANOVA F_2,22_ = 3.644, p = 0.043) (Fig. 2C, Table 1). There was no significant difference in taxonomic richness among the three habitat types (permutational ANOVA F_2,22_ = 2.644, p = 0.094; Fig. 2B, Table 1).

**Figure 2.**
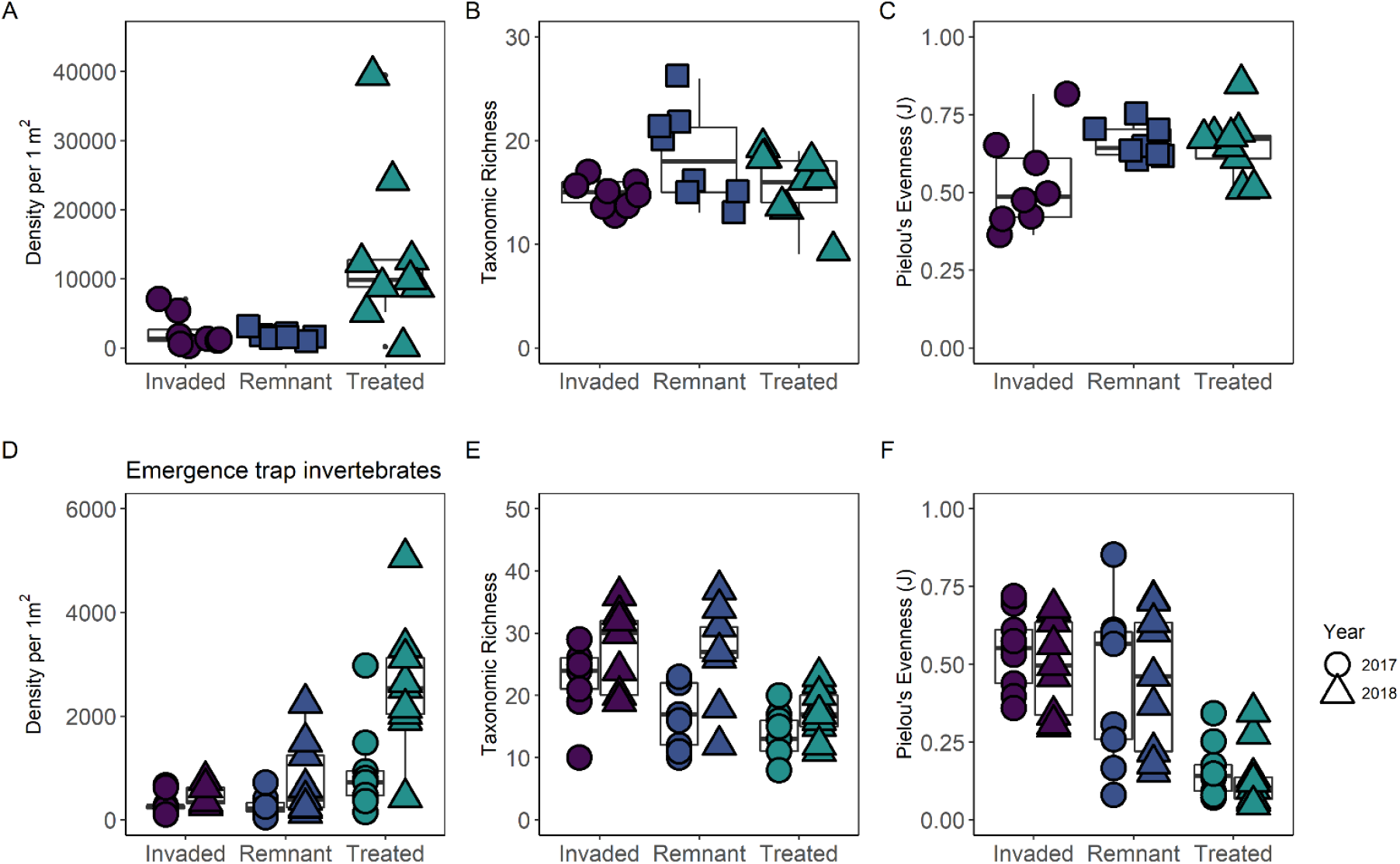
Differences among the three habitat types in aquatic sample macroinvertebrate density per 1 m^2^ (A), taxonomic richness (B), and Pielou’s evenness (C) and emergence trap macroinvertebrate density per m^2^ (D), taxonomic richness (E), and Pielou’s evenness (F). Boxplot whiskers represent 1.5 * IQR / sqrt(n), and notches represent 25%, 50% and 75% quantiles. Aquatic samples include taxa that lacked a winged adult stage while emergence trap samples includes terrestrial, semi-aquatic, and aquatic taxa.

**Table 1.** Average taxonomic richness, invertebrate density, and Pielou’s evenness for aquatic invertebrates in 2018. Samples included taxa that lacked a winged adult stage. Values in parenthesis represent standard error.

|  | Remnant marsh | *P. australis*-invaded sites | Herbicide-treated sites |
| --- | --- | --- | --- |
| Taxonomic richness | 18.50 (± 1.57) | 15.00 (± 0.46) | 15.67 (± 1.07) |
| Density per 1 m<sup>2</sup> | 1176.0 (± 221.92) | 2335.50 (± 879.86) | 13497.33 (± 3892.91) |
| Pielou's evenness | 0.66 (± 0.02) | 0.53 (± 0.05) | 0.65 (± 0.03) |

#### Emergence trap invertebrates

In 2017, we detected 64 taxa in *P. australis*, 36 taxa in herbicide-treated sites, and 53 taxa in remnant marsh (Table S6). Invertebrate densities were significantly higher in herbicide-treated than in *P. australis*-invaded habitat or remnant uninvaded habitat (permutational ANOVA F_2,24_ = 4.521, p = 0.022; Fig. 2D, Table 2). Taxonomic richness was significantly higher in *P. australis*-invaded habitat compared to herbicide-treated and remnant marsh habitat (permutational ANOVA F_2,24_ = 7.705, p = 0.003; Fig. 2E, Table 2) and Pielou’s evenness was significantly lower in the herbicide-treated sites than in the other vegetation types (permutational ANOVA F_2,24_ = 12.119, p < 0.001; Fig 2F, Table 2).

**Table 2.** Average taxonomic richness, invertebrate density, and Pielou’s evenness for invertebrates collected in emergence traps in 2017 and 2018. Emergence traps were sampled from June 19^th^ to July 21^st^ in 2017 and May 20^th^ to July 23^rd^ in 2018. Values in parenthesis represent standard error.

|  | Remnant marsh |  | *P. australis* -invaded sites |  | Herbicide-treated sites |  |
| --- | --- | --- | --- | --- | --- | --- |
|  | 2017 | 2018 | 2017 | 2018 | 2017 | 2018 |
| Taxonomic richness | 16.78<br>(± 1.70) | 26.44<br>(± 2.56) | 22.33<br>(± 1.86) | 27.11<br>(± 2.14) | 13.22<br>(± 1.37) | 17.11<br>(± 1.38) |
| Density per 1 m<sup>2</sup> | 268.33<br>(± 67.21) | 761.44<br>(± 246.92) | 316.56<br>(± 65.22) | 449.00<br>(± 67.21) | 932.22<br>(± 286.62) | 2580.44<br>(± 417.38) |
| Pielou's evenness | 0.45<br>(± 0.08) | 0.45<br>(± 0.08) | 0.54<br>(± 0.04) | 0.49<br>(± 0.05) | 0.16<br>(± 0.03) | 0.14<br>(± 0.04) |

In 2018, we detected a total of 68 taxa in *P. australis* stands, 43 taxa in herbicide-treated sites, and 66 taxa in remnant marsh (Table S6). Invertebrate densities were significantly higher in herbicide-treated sites compared to the other habitat types (permutational ANOVA F_2,24_ = 16.582, p < 0.001; Fig. 2D, Table 2). Taxonomic richness was significantly lower in herbicide-treated sites, while *P. australis*-invaded sites and remnant marsh were similar (permutational ANOVA F_2,24_ = 7.198, p = 0.004; Fig. 2E, Table 2). As in 2017, Pielou’s evenness was significantly lower in herbicide-treated sites compared to *P. australis-*invaded sites or remnant marsh (permutational ANOVA F_2,24_ = 11.914, p < 0.001; Fig. 2F, Table 2).

### Multivariate

#### Aquatic invertebrates

The mean distance to the group centroid was 0.447 in *P. australis-* invaded sites, 0.454 in herbicide-treated sites, and 0.486 in remnant marsh sites, indicating beta diversity of aquatic macroinvertebrates was slightly higher in remnant marsh. However, there was no significant difference in the homogeneity of variances among the habitat types (PERMDISP, F_2,24_ = 0.330, p = 0.766). Because aquatic macroinvertebrates did not differ in the homogeneity of variances among habitat types, we interpret the statistically significant perMANOVA (pseudo-F_2,24_ = 4.379, p = 0.001) as indicating a difference in community location, rather than community dispersion (Warton et al. 2012). More, each habitat type was significantly different from the other (Bonferroni corrected p-value < 0.01 for each comparison).

The final NMDS ordination solution was 3-dimensional, with an acceptable final stress of 0.145, and two convergent solutions found after 28 tries, with a non-metric r^2^ of 0.979 (Fig. 3). On axis 1 and 2 there was little overlap of the three habitat types. Remnant marsh and herbicide-treated sites separated along axis 1, while *P. australis*-invaded sites and herbicide-treated sites separated along axis 2. Remnant marsh was more diverse than either *P. australis* or herbicide-treated sites. Chironomidae (39,024 individuals per 1 m^2^) were associated with herbicide-treated sites and were highly abundant compared to remnant (2,176 individuals per 1 m^2^) or *P. australis*-invaded sites (2,024 individuals per 1 m^2^) sites. Caenidae (small square-gill mayflies) and Leptoceridae (long-horned caddisflies) were also associated with herbicide-treated sites, as were Hydrozoa (hydrozoans) and Gastropoda (snails). Remnant marsh was characterized by Staphylinidae (rove beetles), Collembola (spring tails), Aranea (spiders), and Hydrophilidae (water scavenger beetles). On axis 1 and 3, herbicide-treated and *P. australis* stands overlapped considerably and had fewer associated taxa than remnant habitat. A few remnant marsh and *P. australis-invaded* marsh shared higher densities of Curculionidae (true weevils) and Coenagrionidae (narrow-winged damselflies) which were orthogonal to sites characterized by Dolichopodidae (long-legged flies). The taxa strongly associated with remnant marsh habitat on axis 1 were also associated with vegetation litter.

**Figure 3.**
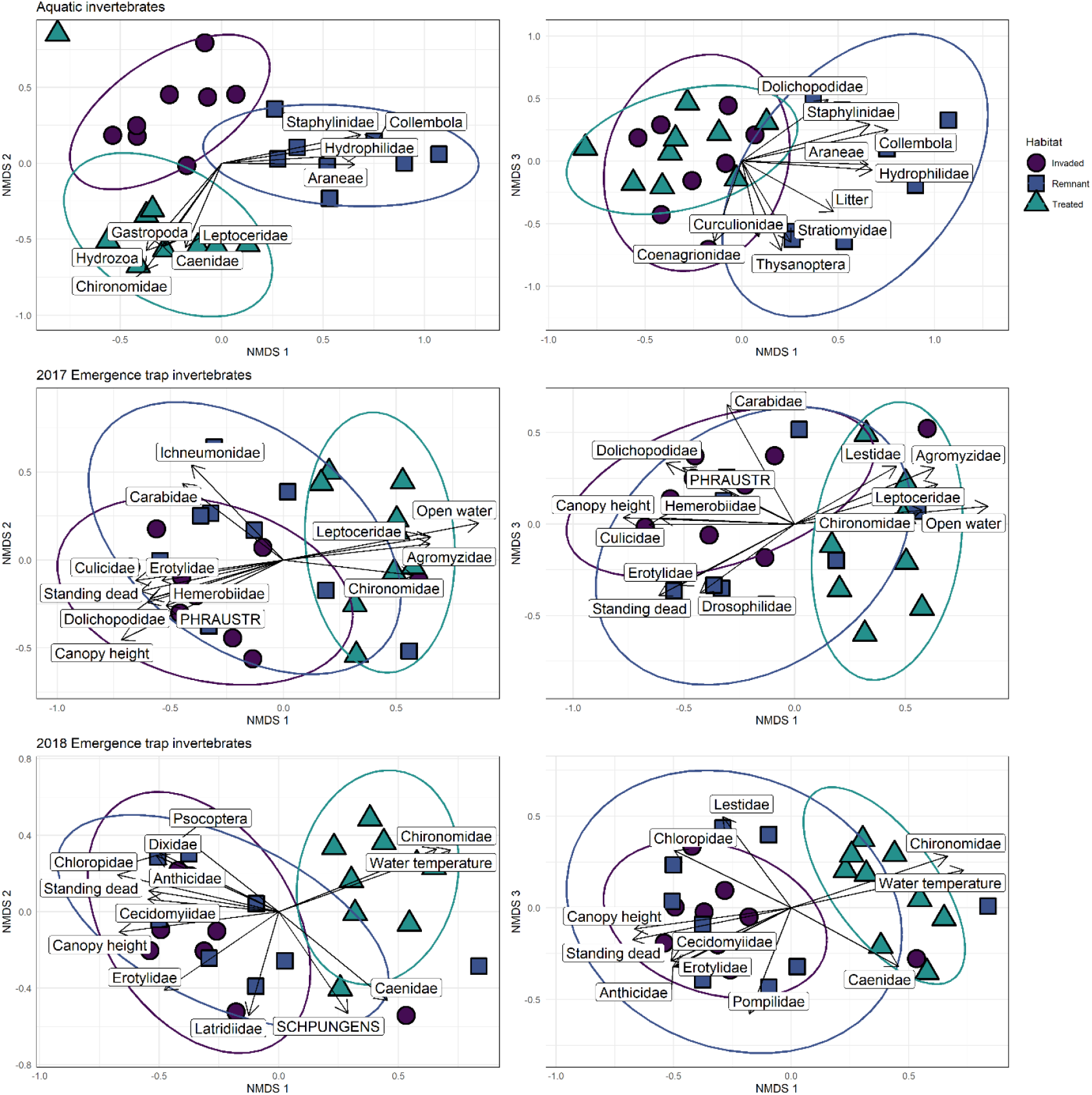
NMDS ordination solutions of aquatic invertebrates and emergence trap invertebrates with reasonably correlated environmental characteristics, plant species, and invertebrate taxa (r^2^ > 0.30) included as vectors and 90% CI ellipses. Plant species codes are presented in all caps and include *Phragmites australis* (PHRAUSTR) and *Schoenoplectus pungens* (SCHPUNGENS), and “standing dead” refers to standing dead plant biomass.

### Emergence trap invertebrates

#### 2017 emergence trap collections

The mean distance to the group centroid was similar among the three habitat types (PERMDISP F_2,24_ = 0.153, p = 0.859), with a mean distance of 0.552 in remnant marsh, 0.539 in *P. australis-invaded* sites and 0.529 in herbicide-treated sites. The significant perMANOVA (pseudo-F_2,24_ = 2.230, p = 0.001) thus indicated a significant difference in the invertebrate communities associated with each habitat type. The invertebrate community in *P. australis*-invaded sites and remnant marsh were not significantly different (Bonferroni corrected p-value = 0.207), while the invertebrate community associated with the herbicide-treated sites was significantly different than *P. australis-invaded* sites (Bonferroni corrected p-value = 0.003) and remnant marsh (Bonferroni corrected p-value = 0.006).

The results of the multivariate generalized linear model indicate that invertebrate community composition among the different habitat types differed significantly over the season, with high densities in herbicide-treated sites that were driven by high densities of Chironomidae over the season (Table 3, Fig. 4A).

**Figure 4.**
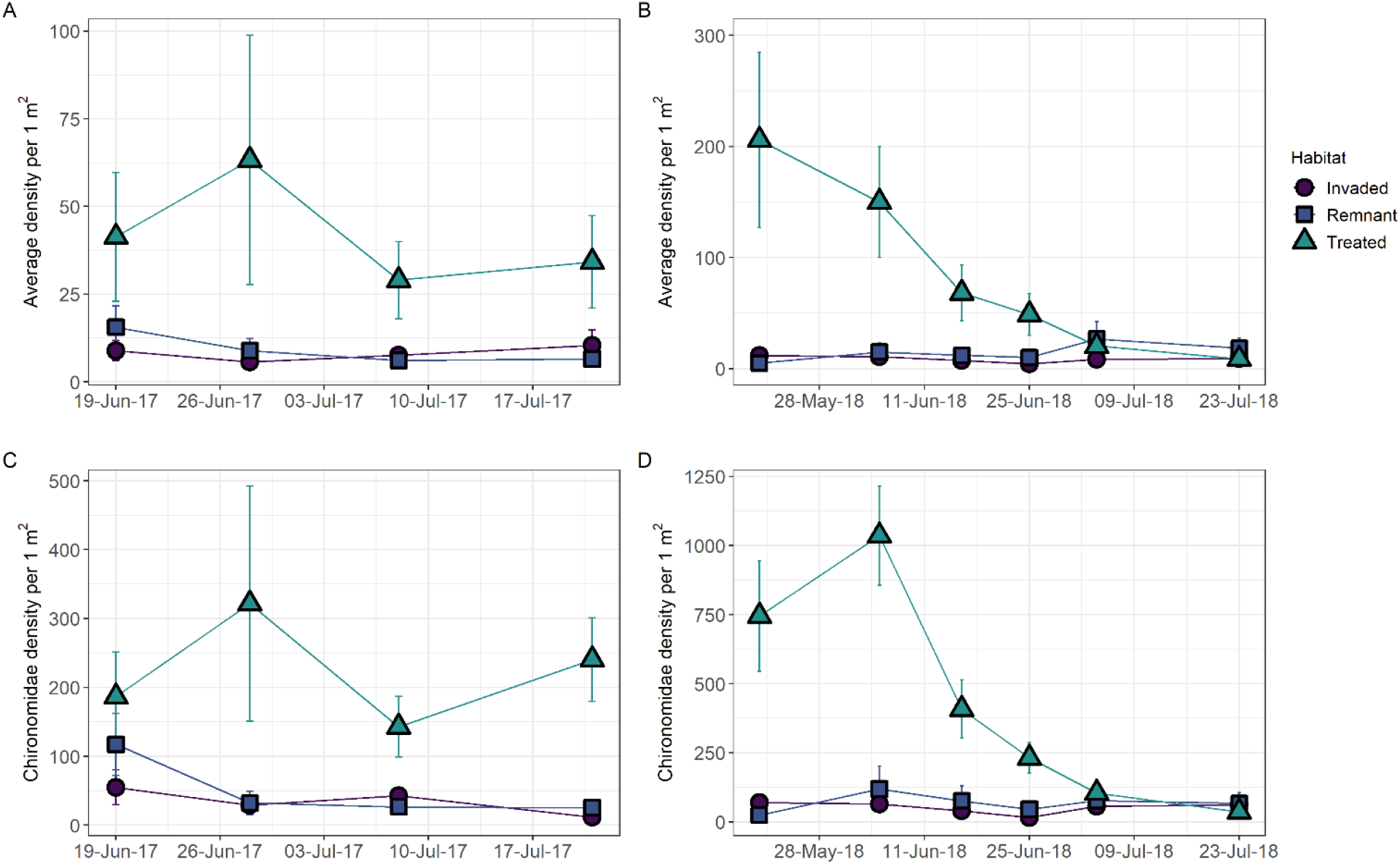
Overall density of invertebrates captured in emergence traps in each habitat type over the field season in 2017 (A) and 2018 (B). Chironomidae densities differed significantly among habitat types over the field season, with high densities in herbicide treated sites in 2017 (C) and 2018 (D). Error bars represent standard error.

**Table 3.** Summary of multivariate generalized linear models of the invertebrate community in 2017 (19-June to 21-July) and 2018 (20-May to 23-July). In both years, macroinvertebrate communities captured in emergence traps differed among the habitat types depending on the time of the season.

|  | 2017 |  |  | 2018 |  |  |
| --- | --- | --- | --- | --- | --- | --- |
|  | Residual df | Deviance | p-value | Residual df | Deviance | p-value |
| Habitat type | 100 | 487.7 | 0.001 | 153 | 679.1 | 0.001 |
| Days | 99 | 225.1 | 0.001 | 152 | 551.6 | 0.001 |
| Habitat type: Days | 97 | 158.9 | 0.001 | 150 | 214.9 | 0.001 |

The final NMDS ordination had a 3D solution with an acceptable final stress of 0.172 after 121 iterations, with non-metric r^2^ of 0.970 (Fig. 3). There was high overlap between *P. australis-*sites and remnant marsh on NMDS axis 1, and herbicide-treated sites were separate. Herbicide-treated sites were associated with high abundances of Chironomidae, Leptoceridae, and Agromyzidae (leaf-miner flies) and characterized by a high percentage of open water. There was high overlap between *P. australis*-invaded sites and remnant marsh sites with high canopy height and more standing dead plant biomass and Culicidae (mosquitos), Erotylidae (pleasing fungus beetles), Hemerobiidae (brown lacewings), and Dolichopodidae (long-legged flies) were all associated with these sites. Ichneumonidae (Ichneumon wasps) and Carabidae (ground beetles) abundances were associated with remnant marsh vegetation with lower canopies and less open water on axis 2. On axis 3, Drosophilidae (small fruit flies) abundances were associated with sites that had a higher percent cover of standing dead plant biomass.

#### 2018 emergence trap collection

There was no significant difference in the homogeneity of variances among the habitat types (PERMDISP F_2,24_ = 0.8156, p = 0.444). The mean distance to the group centroid was 0.506 in *P. australis-invaded* sites, 0.492 in herbicide-treated sites, and 0.541 in remnant marsh sites. The invertebrate communities were significantly different among the three habitat types (perMANOVA pseudo-F_2,24_ = 2.941, p = 0.001). The invertebrate community in remnant marsh and *P. australis*-invaded sites were not significantly different from one another (Bonferroni corrected p-value = 0.067), while the invertebrate community in herbicide-treated sites was significantly different from the community in *P. australis-*invaded sites (Bonferroni corrected p-value = 0.002) and from the community in remnant marsh sites (Bonferroni corrected p-value = 0.001).

The results of the multivariate generalized linear model reveal that the invertebrate community composition among the different habitat types differed significantly over the field season (Table 3, Fig. 4B). Herbicide-treated sites had higher numbers of invertebrates, driven by Chironomidae, which had very high abundances at the beginning of the season (Fig. 4D).

The final NMDS ordination was a 3D solution with a stress of 0.159 after 81 iterations and a non-metric r^2^ of 0.975 (Fig. 3). There was considerable overlap in the invertebrate community composition between *P. australis* and remnant marsh on all three axes, while herbicide-treated sites were separated from the other habitats along axis 1. Chironomidae (20,809 individuals per m^2^) were highly abundant in herbicide-treated sites compared to remnant marsh (3,472 individuals per m^2^) and *P. australis* (2,621 individuals per m^2^) sites. Caenidae were also associated with herbicide-treated sites that had higher *Schoenoplectus pungens* (common three-square rush [(Vahl) Palla]) cover. Remnant marsh and *P. australis-invaded* sites had similar community composition, and higher canopy cover and standing dead as a result of their abundant emergent vegetation. Psocoptera (booklice), Anthicidae (ant beetles) and Diptera such as Dixidae, Chloropidae, and Cecidomyiidae were associated with remnant marsh or *P. australis* stands with slightly more standing dead litter on axis 2, while Erotylidae were associated with canopy height. On axis 3, there is a separation between remnant marsh and *P. australis* stands that support Lestidae (spread-winged damselflies) and those that support Pompilidae (spider-wasps), a parasitic wasp whose host taxa include *Dolomedes* (fishing spiders) (Merritt et al. 2008).

#### Congruence between aquatic and emerging invertebrates

When we assessed the congruence between the subset of invertebrates that had an aquatic stage and an emergent terrestrial stage, the invertebrate communities were significantly congruent with relatively high t-values in both remnant marsh (Procrustes analysis; t = 0.865, p = 0.049) and *P. australis-invaded* marsh (Procrustes analysis; t = 0.894, p = 0.006). In this reduced subset of data, *P. australis* and remnant marsh each had an emerging invertebrate taxonomic richness of 28, and Chironomidae was the most abundant in both habitat types. The taxonomic richness of *P. australis* aquatic invertebrates was lower (10 taxa) than remnant marsh (17 taxa).

The aquatic and emerging invertebrate communities in herbicide-treated habitat had lower, non-significant congruence (Procrustes analysis; t = 0.485, p = 0.647). In the herbicide-treated sites, there were 17 taxa present in the aquatic samples and Chironomidae, Ceratopogonidae, Caenidae and Leptoceridae had the highest abundances. Of the emergence trap samples, there were 24 taxa present, and Chironomidae and Ceratopogonidae were the overwhelming majority. There were 388 Leptoceridae per 1 m^2^ in aquatic samples and 195 per 1 m^2^ in the emergence trap samples in these sites, and 8,560 Caenidae per 1 m^2^ in the aquatic samples and only 17 individuals per 1 m^2^ in the emergence trap samples in herbicide treated sites. NMDS ordinations visualizing aquatic and emerging invertebrate communities by habitat type are presented in Figure S2.

## Discussion

Our results suggest that *P. australis* invasion does have an influence on the macroinvertebrate community occupying aquatic habitat, but the drastic change to vegetation caused by the herbicide-based control of *P. australis* in freshwater marshes has a profound influence on the macroinvertebrate community and may select for tolerant taxa that feed on decomposing organic matter and periphyton such as Chironomidae and Caenidae, or that may prey on these consumers, such as Leptoceridae.

We sought to address three knowledge gaps regarding the effects of *P. australis* invasion and herbicide-based control on the macroinvertebrate community in freshwater coastal marsh. First, we asked whether macroinvertebrate communities would be less dense and less diverse in *P. australis-*invaded marsh than in remnant, uninvaded marsh. Though evenness was lower in *P. australis*-invaded marsh, taxonomic richness and densities did not differ for the aquatic macroinvertebrates. In terms of macroinvertebrates sampled with emergence traps, we found no significant differences between the density or evenness of the *P. australis-*invaded sites and remnant, uninvaded marsh habitats, and richness in 2017 was actually higher in the *P. australis*-invaded sites. However, this difference was not evident in 2018, when we were able to capture early-season emergence. Our findings contrast with work from the southern shore of Lake Erie that indicated benthic invertebrate diversity increased with *P. australis* cover, though our results do agree with their finding that densities were similar between *P. australis* and uninvaded marsh (Holomuzki & Klarer 2010). Another study in Lake Erie marshes also found higher densities of aquatic macroinvertebrates in *P. australis* compared to *Typha*, which was a result of gastropod and chironomid abundance (Kulesza et al. 2008). In our study, the total densities for both chironomids and gastropods were similar between *P. australis* and remnant uninvaded marsh. Despite similarities in richness, evenness, and density the aquatic macroinvertebrate community composition was significantly different between *P. australis* and remnant marsh. Numerous aquatic (e.g., Hydrophilidae, Stratiomyidae) and semi-aquatic (e.g., Staphylinidae, Collembola, Dolichopodidae) taxa had their densities strongly, positively associated with remnant marsh whereas none of the taxa densities were strongly positively correlated with *P. australis-invaded* sites. The varying water depths and diverse vegetation in remnant marsh may provide more aquatic habitat diversity for macroinvertebrates than the less diverse *P. australis* habitat. Similar patterns were observed in North American brackish marshes, where native vegetation provided more refuge for macroinvertebrates than *P. australis* (Angradi et al. 2001).

Second, we asked whether the macroinvertebrate community in herbicide-treatment areas more closely resembled remnant, uninvaded marsh or *P. australis-*invaded marsh. We found that treated marsh was dissimilar to both. Herbicide-treated sites supported higher densities in both the aquatic and emergence trap samples, and lower richness and evenness in the emergence traps samples than either remnant or *P. australis*-invaded habitats. This was principally due to dominance by Diptera, particularly high abundance of Chironomidae. The herbicide treated sites also had higher densities of Leptoceridae and Caenidae in both aquatic and emergence trap samples and the aquatic samples had higher densities of Gastropoda and Hydrozoa. All of these are taxa that likely benefited from the herbicide-caused die-off of dense *P. australis* plants, the resulting increased in submerged litter (Yuckin 2018) and the submerged and floating aquatic vegetation that dominated these sites for at least two-years after their treatment (Robichaud & Rooney 2021a). Chironomids, which were extremely abundant in herbicide-treated plots, are fast-growing and opportunistic, and experiments have demonstrated they will preferentially select sites with high food quality in organically enriched ecosystems (de Haas et al. 2006). Interestingly, there exist studies on the effects of glyphosate application directly to wetlands and these report similar community changes. E.g., after a glyphosate-based herbicide was directly applied to experimental wetlands in New Brunswick, chironomid abundances increased significantly following macrophyte die-offs (Baker et al. 2014). In the Prairie Pothole Region, the removal of *Typha* spp. using a glyphosate-based herbicide led to higher abundances of Corixidae and Chironomidae (Linz et al. 1999). Macroinvertebrate densities were higher in *P. australis* and herbicide-treated *P. australis*, compared to *Typha*, in marshes on the southern coast of Lake Erie, due in part to high abundances of chironomids (Kulesza et al. 2008). Promisingly, the removal of *P. australis* in brackish marshes resulted in macroinvertebrate communities that resembled native vegetation within five years of treatment (Gratton & Denno 2005).

*Phragmites australis* and remnant marsh vegetation both supported aquatic, semi-aquatic, and terrestrial taxa including multiple Families of Coleoptera (Anthicidae, Erotylidae, Latridiidae) and Diptera (Dixidae, Cecidomyiidae, Chloropidae). Few studies have compared invertebrate communities using emergence traps among *P. australis* and native vegetation, and fewer still have done so in freshwater marshes. In a brackish *Spartina* marsh (New Jersey, USA) Gratton & Denno (2005) found that the macroinvertebrate community in *P. australis* was significantly different than the community in *Spartina*, and was characterized by Collembola, chironomids, and a reduction in marsh spiders due to the structural changes of invasion. In the wetlands of the Great Salt Lake (Utah, USA), *P. australis* provided adequate habitat for both aquatic and terrestrial arthropods and supported similar assemblages as native hardstem bulrush (*Schoenoplectus acutus*) and alkali bulrush (*Bolboshoenus maritimus*) habitat (Leonard et al. 2021). However, native pickleweed (*Salicornia rubra*) had a significantly different arthropod assemblage than the other three vegetation types which is attributed to this vegetation community having less water, litter, and vegetation biomass (Leonard et al. 2021). The similarities in emergence trap macroinvertebrate communities between *P. australis* and remnant marsh in our study also indicate that habitat characteristics, such as plant biomass and litter, are important determinants of the emerging macroinvertebrate communities. Particularly, much of the remnant marsh habitat in our study was dominated by *Typha*, likely the invasive hybrid *Typha* x *glauca*, which creates tall dense monocultures similar to *P. australis*.

Our third question asked whether the community congruence between the subset of taxa that have an aquatic juvenile stage and a winged adult stage would be greater in remnant marsh than in *P. australis*-invaded sites or herbicide-treated sites. Though the invertebrate communities in herbicide-treated sites had some of the same dominant taxa, they exhibited less congruence than the invertebrate communities in the *P. australis*-invaded sites and in the remnant, uninvaded marsh sites. In contrast, the aquatic and emerging macroinvertebrate communities were significantly congruent in the *P. australis*-invaded sites and remnant sites. This contradicted our prediction that *P. australis* invasion would disrupt the agreement among aquatic and emerging macroinvertebrate communities and was surprising given that the emergence trap collections in *P. australis*-invaded sites and remnant marsh sites were indistinguishable, whereas the aquatic sample collections from these two habitat types were distinctive. We expect this is because of the abundant above-ground biomass in both vegetation communities that provide ample substrate for emerging macroinvertebrates.

The lack of emergent vegetation in herbicide-treated sites likely limited the ability of certain taxa to emerge and selects against taxa, such as Cecidomyiidae, that require vegetation for a portion of their life cycle. Some herbicide-treated sites were characterized by higher densities of Caenidae (Ephemeroptera). These same sites had higher abundances of emergent vegetation (i.e., *Schoenoplectus pungens*), which likely served the important role of providing a substrate that aquatic nymphs can use to crawl out of the water and molt. In herbicide-treated sites with less vegetation, the sparse emergent canopy led to warmer water temperatures and high densities of chironomids were positively associated with these sites. As chironomids can float to the surface of the water and emerge, they are not limited by a lack of emergent vegetation or substrate the way other invertebrate taxa are. In general, warm water temperatures and high-quality food correlates with shorter life cycles in chironomids (Merritt et al. 2008) which may explain why we saw such high densities emerging in these sites. The differences in macroinvertebrate community and congruence may not be entirely attributed to emergent vegetation. We expect the shallow open-water areas that are created after herbicide-treatment may provide better foraging grounds for insectivorous fish, birds, and other predators that could selectively remove certain macroinvertebrates. Further, we only sampled aquatic macroinvertebrates at one time point (early May) to capture the taxa present before emergence. We suggest that future research sample this community multiple times over the season to characterize communities more accurately in different habitat.

Our work took place alongside a wetland restoration project initiated by government partners, and as such there are constraints to our study design. We were unable to collect samples from before herbicide treatment began, thus limiting our ability to compare macroinvertebrate communities to a sufficient baseline. However, the effect size we present is large and consistent across two consecutive years – there are significantly more macroinvertebrates emerging from herbicide-treated sites one- and two-years after treatment compared to other vegetation communities in the marsh. Additionally, these results align with similar work that noted high densities of invertebrates following glyphosate-based herbicide treatment (e.g., Linz et al. 1999, Baker et al. 2014) or mechanical disturbance (e.g., Schummer et al. 2012). Finally, while we do not have macroinvertebrate community data from sites before they were treated, we do have extensive monitoring results of the environmental variables and vegetation community at sites before and after treatment in Long Point, our study site, and Rondeau Provincial Park, another wetland complex on Lake Erie. Aerial application of a glyphosate-based herbicide consistently resulted in a significant and dramatic alteration to marsh vegetation (Robichaud and Rooney 2021a). While the treated sites in 2016 were close to one another, the *P. australis* stands we sampled in 2017 and 2018 were dense monocultures that were equivalent to what was treated in 2016 and have since been treated in a similar manner. Thus, while we do not have ‘before’ samples we are confident in our conclusion that the treatment of *P. australis* stands and the subsequent change in wetland vegetation from dense emergent vegetation to shallow open water wetland with sparse vegetation resulted in a change in the macroinvertebrate community.

Comparing the way native biota use invaded, uninvaded, and herbicide-treated habitat can inform land management by weighing the consequences of unmitigated invasion against any potential unintended impacts of invasive species control. In the case of *P. australis*, the effects of invasion potentially include altering the aquatic macroinvertebrate community composition. However, our results also emphasize that efforts to control invasive plant species can have significant effects on native invertebrate communities that last beyond the initial treatment. Measuring these changes in the macroinvertebrate community is important as macroinvertebrates are key components of aquatic food webs and aquatic-terrestrial linkages (e.g., Collier et al. 2002). We recommend that restoration projects include some monitoring of the macroinvertebrate community, and that wetland-related projects sample the aquatic, semi-aquatic, and terrestrial macroinvertebrate communities to holistically assess community responses. Given the importance of a diverse macroinvertebrate community, we emphasize that for restoration to be successful, in the macroinvertebrate community composition of herbicide-treated areas should eventually converge with that found in remnant, uninvaded marsh. Thus, longer term monitoring is necessary to assess restoration success.

## Acknowledgements

We would like to thank the research technicians and graduate students who assisted with field and lab work on this project: Jessie Pearson, Sarah Yuckin, Heather Polowyk, Calvin Lei, Laura Beecraft, Matthew Bolding, Hillary Quinn-Austin, Bailey Ruest, and Neeva Demules. Thank you to Heidi Swanson, Claude Lavoie, and Marcel Pinheiro for comments on an early draft of the manuscript. Special thanks to Jennifer Gleason for assistance with invertebrate ecology, and to two anonymous reviewers who greatly improved the manuscript. This project was supported by funding from NSERC PGS (Robichaud) and from the Species at Risk Research Fund of Ontario (RF_35_18_UWater2) granted to Robichaud and Rooney.

## Data Availability Statement

Data and scripts used in analyses for this paper are available in a public GitHub repository (https://github.com/cdrobich/macroinvert_phragmites.git).

## Notes

https://github.com/cdrobich/macroinvert_phragmites.git

### Competing Interest Statement

The authors have declared no competing interest.

## References

Anderson MJ (2006) Distance-based tests for homogeneity of multivariate dispersions. Biometrics 245–253

Anderson MJ, Ellingsen KE, McArdle BH (2006) Multivariate dispersion as a measure of beta diversity. Ecology Letters 9:683–693

Anderson J.T. & Davis C.A. (2013). Wetland Techniques, Volume 2: Organisms. (Ed. Springer), New York, NY.

Angradi TR, Hagan SM, Able KW (2001) Vegetation type and the intertidal macroinvertebrate fauna of a brackish marsh: *Phragmites* vs. *Spartina*. Wetlands 21:75–92

Baker LF et al. (2014) The direct and indirect effects of a glyphosate-based herbicide and nutrients on Chironomidae (Diptera) emerging from small wetlands. Environmental Toxicology and Chemistry 33:2076–2085

Ball H et al. (2003) The Ontario Great Lakes Coastal Weltand Atlas: A summary of information (1983-1997). Environment Canada 49

Batzer DP (2013) The seemingly intractable ecological responses of invertebrates in North American Wetlands: A review. Wetlands 33:1–15

Batzer DP, Rader RB, Wissinger SA (1999) Invertebrates in freshwater wetlands of North America: ecology and management. John Wiley & Sons, Inc.

Batzer DP, Wu H (2020) Ecology of terrestrial arthropods in freshwater wetlands. Annual Review of Entomology 65:101–119

Bauer JT (2012) Invasive species: ‘back-seat drivers’ of ecosystem change? Biological Invasions 14:1295–1304

Bonada N et al. (2006) Developments in aquatic insect biomonitoring: A comparative analysis of recent approaches. Annual Review of Entomology 51:495–523

Brinson MM, Malvárez AI (2021) Temperate freshwater wetlands: types, status, and threats. Environmental Conseration 29:115–133

Collier K., Bury S, Gibbs M (2002) A stable isotope study of linkages between stream and terrestrial food webs through spider predation. Freshwater Biology 47:1651–1659

Crooks JA (2002) Characterizing ecosystem-level consequences of biological invasions: the role of ecosystem engineers. Oikos 97:153–166

D’Antonio CM, Meyerson LA (2002) Exotic plant species as problems and solutions in ecological restoration. Restoration Ecology 10:703–713

Fell PE et al. (2006) Short-term effects on macroinvertebrates and fishes of herbiciding and mowing *Phragmites australis*-dominated tidal marsh. Northeastern Naturalist 13:191–212

Fruh D, Stoll S, Haase P (2012) Physicochemical and morphological degradation of stream and river habitats increases invasion risk. Biological Invasions 14:2243–2253

Gathman JP, Burton TM (2011) A Great Lakes coastal wetland invertebrate community gradient: Relative influence of flooding regime and vegetation zonation. Wetlands 31:329–341

Gratton C, Denno RF (2005) Restoration of arthropod assemblages in a spartina salt marsh following removal of the invasive plant *Phragmites australis*. Restoration Ecology 13:358–372

Greenberg DA, Green DM (2013) Effects of an invasive plant on population dynamics in toads. Conservation Biology 27:1049–1057

de Haas E et al. (2006) Habitat selection by chironomid larvae: fast growth. Journal of Animal Ecology 148–155

Harvey KJ, Britton DR, Minchinton TE (2014) Detecting impacts of non-native species on associated invertebrate assemblages depends on microhabitat. Austral Ecology 39:511–521

Hazelton ELG et al. (2014) *Phragmites australis* management in the United States: 40 years of methods and outcomes. AoB PLANTS 6:1–19

Herdendorf CE (1992) Lake Erie coastal wetlands: an overview. Journal of Great Lakes Research 18:533–551

Holomuzki JR, Klarer DM (2010) Invasive reed effects on benthic community structure in Lake Erie coastal marshes. Wetlands Ecology and Management 18:219–231

Jung JA, Rokitnicki-Wojcik D, Midwood JD (2017) Characterizing past and modelling future spread of *Phragmites australis* ssp*. australis* at Long Point Peninsula, Ontario, Canada. Wetlands 37:961–973

Kulesza AE, Holomuzki JR, Klarer DM (2008) Benthic community structure in stands of *Typha angustifolia* and herbicide – treated and untreated *Phragmites australis*. Wetlands 28:40–56

Leonard EE et al. (2021) Arthropod assemblages in invasive and native vegetation of Great Salt Lake Wetlands. Wetlands 41:1–17

Linz GM et al. (1999) Response of invertebrates to glyphosate-induced habitat alterations in wetlands. Wetlands 19:220–227

MacDougall A, Turkington R (2005) Are invasive species the drivers or passengers of change in degraded ecosystems? Ecology 86:42–55

Le Maitre DC (2004) Predicting invasive species impacts on hydrological processes: the consequences of plant physiology for landscape processes. Weed Technology 18:1408–1410

Markle CE, Chow-Fraser P (2018) Effects of European common reed on Blanding’s turtle spatial ecology. Journal of Wildlife Management 82:857–864

Marshall SA (2006) Insects: Their Natural History and Diversity: with a Photographic Guide to Insects of Eastern North America. Firefly Books

de Mendiburu F (2020) agricolae: Statistical Procedures for Agricultural Research.

Merritt RW, Cummins KW, Berg MB (2008) An introduction to the aquatic insects of North America. 4th ed. Kendall Hunt Publishing Company, Dubuque, IA

Oksanen J et al. (2020) vegan: Community Ecology Package. R package version 2.5-7.

R Core Team (2020) R: A language and environment for statistical computing. R Foundation for Statistical Computing, Vienna, Austria

Robichaud CD, Rooney RC (2017) Long-term effects of a Phragmites australis invasion on birds in a Lake Erie coastal marsh. Journal of Great Lakes Research 43

Robichaud C.D, Rooney R. (2021a) Effective suppression of established invasive *Phragmites australis* leads to secondary invasion in a coastal marsh. Invasive Plant Science and Management 14:9–19

Robichaud C.D., Rooney RC (2021b) Low concentrations of glyphosate in water and sediment after direct over-water application to control an invasive aquatic plant. Water Research 188

Schummer ML et al. (2012) Comparisons of bird, aquatic macroinvertebrate, and plant communities among dredged ponds and natural wetland habitats at Long Point, Lake Erie, Ontario. Wetlands 32:945–953

Talley TS, Levin LA (2001) Modification of sediments and macrofauna by an invasive marsh plant. Biological Invasions 3:51–68

Thorp JH, Covich AP (1991) Ecology and Classification of North American freshwater invertebrates. Academic Press, Inc., San Diego, California.

Triplehorn CA, Johnson NF (2005) Borror and Delong’s Introduction to the Study of Insects. Thompson Brooks/Cole

Tulbure MG, Johnston CA (2010) Environmental conditions promoting non-native *Phragmites australis* expansion in great lakes coastal wetlands. Wetlands 30:577–587

Vilà M et al. (2011) Ecological impacts of invasive alien plants: a meta-analysis of their effects on species, communities and ecosystems. Ecology letters 14:702–8

Warton DI, Wright ST, Wang Y (2012) Distance-based multivariate analyses confound location and dispersion effects. Methods in Ecology and Evolution 3:89–101

Weidlich EWA et al. (2020) Controlling invasive plant species in ecological restoration: A global review. Journal of Applied Ecology 57:1806–1817

Wheeler B, Torchiano M (2016) lmPerm: Permutation tests for linear models.

Whickham H (2016) ggplot2: Elegant Graphics for Data Analysis.

Wikum DA, Shanholtzer GF (1978) Application of the Braun-Blanquet cover-abundance scale for vegetation analysis in land development studies. Environmental Management 2:323–329

Wilcox DA (2012) Response of wetland vegetation to the post-1986 decrease in Lake St. Clair water levels: Seed-bank emergence and beginnings of the *Phragmites australis* invasion. Journal of Great Lakes Research 38:270–277

Wilcox KL et al. (2003) Historical distribution and abundance of *Phragmites australis* at Long Point, Lake Erie, Ontario. Journal of Great Lakes Research 29:664–680

Yuckin S (2018) Detecting the effects of biological invasion and subsequent control efforts on wetland ecological processes. University of Waterloo. MSc thesis http://hdl.handle.net/10012/13888

Yuckin S, Rooney R (2019) Significant increase in nutrient stocks following *Phragmites australis* invasion of freshwater meadow marsh but not of cattail marsh. Frontiers in Environmental Science 7:1–16

Zedler JB, Kercher S (2004) Causes and consequences of invasive plants in wetlands: Opportunities, opportunists, and outcomes. Critical Reviews in Plant Sciences 23:431–452

