## Supplementary information for "Control of invasive *Phragmites australis* (European common reed) alters macroinvertebrate communities"

Appendix A. Study sites used to evaluate aquatic and emerging invertebrate communities in Long Point, ON. Sites were established along a water-depth gradient and traps were visited approximately every 10 days. Collection vessels were not always successfully retrieved, typically due to weather resulting in the loss of a vessel, and the number of collections per site are noted.

| Site | Habitat type | Year | Vegetation type | Water depth (cm) | Number of collections |
|---|---|---|---|---|---|
| IN1 | Invaded | 2017 | *Phragmites* | 35.5 | 4 |
| IN2 | Invaded | 2017 | *Phragmites* | 28.7 | 4 |
| IN3 | Invaded | 2017 | *Phragmites* | 35.0 | 4 |
| IN4 | Invaded | 2017 | *Phragmites* | 36.5 | 3 |
| IN5 | Invaded | 2017 | *Phragmites* | 22.3 | 4 |
| IN6 | Invaded | 2017 | *Phragmites* | 43.3 | 4 |
| IN7 | Invaded | 2017 | *Phragmites* | 36.7 | 4 |
| IN8 | Invaded | 2017 | *Phragmites* | 65.7 | 4 |
| INOW | Invaded | 2017 | *Phragmites* | 70.0 | 4 |
| RES1 | Herbicide-treated | 2017 | Herbicide-treated | 46.0 | 4 |
| RES2 | Herbicide-treated | 2017 | Herbicide-treated | 40.7 | 2 |
| RES3 | Herbicide-treated | 2017 | Herbicide-treated | 48.8 | 4 |
| RES4 | Herbicide-treated | 2017 | Herbicide-treated | 50.7 | 4 |
| RES5 | Herbicide-treated | 2017 | Herbicide-treated | 32.2 | 4 |
| RES6 | Herbicide-treated | 2017 | Herbicide-treated | 55.5 | 4 |
| RES7 | Herbicide-treated | 2017 | Herbicide-treated | 40.3 | 4 |
| RES8 | Herbicide-treated | 2017 | Herbicide-treated | 66.8 | 4 |
| RESOW | Herbicide-treated | 2017 | Herbicide-treated | 75.0 | 3 |
| UNM1 | Remnant | 2017 | Meadow | 27.3 | 4 |
| UNM2 | Remnant | 2017 | Meadow | 32.3 | 4 |
| UNM3 | Remnant | 2017 | Meadow | 21.2 | 4 |
| UNM4 | Remnant | 2017 | Meadow | 17.0 | 3 |
| UNOW | Remnant | 2017 | Emergent | 75.0 | 4 |
| UNT1 | Remnant | 2017 | Emergent | 37.7 | 4 |
| UNT2 | Remnant | 2017 | Emergent | 36.5 | 4 |
| UNT3 | Remnant | 2017 | Emergent | 47.3 | 4 |
| UNT4 | Remnant | 2017 | Emergent | 44.0 | 4 |
| IN1 | Invaded | 2018 | *Phragmites* | 23.3 | 6 |
| IN2 | Invaded | 2018 | *Phragmites* | 58 | 6 |
| IN3 | Invaded | 2018 | *Phragmites* | 22.3 | 6 |
| IN4 | Invaded | 2018 | *Phragmites* | 28.7 | 6 |
| IN5 | Invaded | 2018 | *Phragmites* | 42 | 6 |

| Site | Habitat type | Year | Vegetation type | Water depth (cm) | Number of collections |
|---|---|---|---|---|---|
| IN6 | Invaded | 2018 | *Phragmites* | 26.3 | 6 |
| IN7 | Invaded | 2018 | *Phragmites* | 70.5 | 6 |
| IN8 | Invaded | 2018 | *Phragmites* | 48.3 | 6 |
| IN9 | Invaded | 2018 | *Phragmites* | 77 | 5 |
| RES1 | Herbicide-treated | 2018 | Herbicide-treated | 26.7 | 6 |
| RES2 | Herbicide-treated | 2018 | Herbicide-treated | 27.2 | 6 |
| RES3 | Herbicide-treated | 2018 | Herbicide-treated | 29.7 | 6 |
| RES4 | Herbicide-treated | 2018 | Herbicide-treated | 35.3 | 5 |
| RES5 | Herbicide-treated | 2018 | Herbicide-treated | 21.7 | 4 |
| RES6 | Herbicide-treated | 2018 | Herbicide-treated | 51 | 6 |
| RES7 | Herbicide-treated | 2018 | Herbicide-treated | 24 | 6 |
| RES8 | Herbicide-treated | 2018 | Herbicide-treated | 23.2 | 6 |
| RES9 | Herbicide-treated | 2018 | Herbicide-treated | 49.7 | 5 |
| UNM1 | Remnant | 2018 | Meadow | 16.3 | 6 |
| UNM2 | Remnant | 2018 | Meadow | 16.7 | 6 |
| UNM3 | Remnant | 2018 | Meadow | 7 | 6 |
| UNM4 | Remnant | 2018 | Meadow | 15 | 6 |
| UNM5 | Remnant | 2018 | Meadow/mixed | 24.7 | 6 |
| UNM6 | Remnant | 2018 | Meadow/mixed | 32.8 | 6 |
| UNT9 | Remnant | 2018 | Emergent | 75 | 5 |
| UNT1 | Remnant | 2018 | Emergent | 34.8 | 6 |
| UNT2 | Remnant | 2018 | Emergent | 36.2 | 6 |

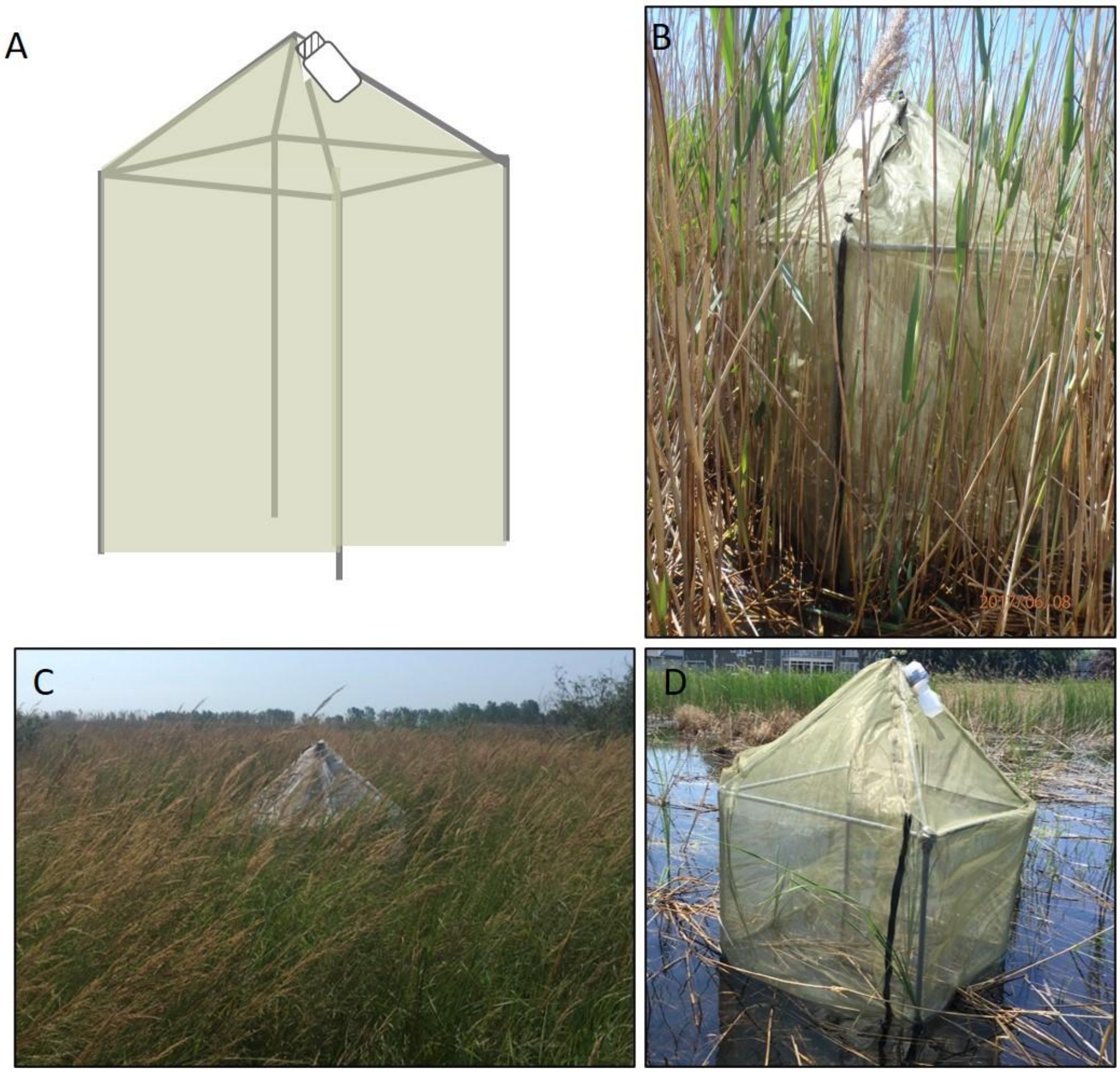


Appendix B. Emergence trap schematic (A), and examples of traps in *P. australis* (B), meadow marsh (C), and herbicide-treated sites (D) in Long Point, ON.

Appendix C. Emerging invertebrate taxa identified in emergence trap (1 $m^2$) samples collected from herbicide-treated, *P. australis*, and remnant marsh sites from Long Point, ON in 2017 and 2018.

| Taxa | Taxonomic level | Order |
|---|---|---|
| Agromyzidae | Subfamily | Diptera |
| Anthicidae | Family | Coleoptera |
| Anthicidae larvae | | |
| Anthocoridae | Family | Hemiptera |
| Anthomyiidae | Family | Diptera |
| Anthomyzidae | Family | Diptera |
| Aphelinidae | Family | Hymenoptera |
| Aphididae | Family | Hemiptera |
| Araneae | Order | |
| Baetidae | Family | Ephemeroptera |
| Braconidae | Family | Hymenoptera |
| Bucculatricidae | Family | Lepidoptera |
| Caenidae | Family | Ephemeroptera |
| Calliphoridae | Family | Diptera |
| Carabidae | Family | Coleoptera |
| Carnidae | Family | Diptera |
| Cecidomyiidae | Family | Diptera |
| Ceraphronidae | Family | Hymenoptera |
| Ceratopogonidae | Family | Diptera |
| Chamaemyiidae | Family | Diptera |
| Chaoboridae | Family | Diptera |
| Chironomidae | Family | Diptera |
| Chloropidae | Family | Diptera |
| Chrysomelidae | Family | Coleoptera |
| Cicadellidae | Family | Hemiptera |
| Coccinellidae | Family | Coleoptera |
| Coccinellidae larvae | | |
| Coccoidea | Superfamily | Hemiptera |
| Coenagrionidae | Family | Odonata |
| Collembola | Order | |
| Cosmopterigidae | Family | Lepidoptera |
| Crambidae | Family | Lepidoptera |
| Cryptophagidae | Family | Coleoptera |
| Culicidae | Family | Diptera |
| Curculionidae | Family | Coleoptera |

| Taxa | Taxonomic level | Order |
|---|---|---|
| Diapriidae | Family | Hymenoptera |
| Dixidae | Family | Diptera |
| Dolichopodidae | Family | Diptera |
| Drosophilidae | Family | Diptera |
| Empididae | Family | Diptera |
| Encyrtidae | Family | Hymenoptera |
| Ephemerellidae | Family | Ephemeroptera |
| Ephydridae | Family | Diptera |
| Erotylidae | Family | Coleoptera |
| Erotylidae larvae | | |
| Eulophidae | Family | Hymenoptera |
| Eurytomidae | Family | Hymenoptera |
| Figitidae | Family | Hymenoptera |
| Formicidae | Family | Hymenoptera |
| Gelechiidae | Family | Lepidoptera |
| Gracillariidae | Family | Lepidoptera |
| Halictidae | Family | Hymenoptera |
| Hemerobiidae | Family | Neuroptera |
| Hydrophilidae | Family | Coleoptera |
| Hydroptilidae | Family | Trichoptera |
| Ichneumonidae | Family | Hymenoptera |
| Lampyridae | Family | Coleoptera |
| Latridiidae | Family | Coleoptera |
| Leptoceridae | Family | Trichoptera |
| Lestidae | Family | Odonata |
| Libellulidae | Family | Odonata |
| Limnephilidae | Family | Trichoptera |
| Megaspilidae | Family | Hymenoptera |
| Miridae | Family | Hemiptera |
| Molannidae | Family | Trichoptera |
| Muscidae | Family | Diptera |
| Mycetophilidae | Family | Diptera |
| Mymaridae | Family | Hymenoptera |
| Nitidulidae | Family | Coleoptera |
| Noctuidae | Family | Lepidoptera |
| Phalacridae | Family | Coleoptera |
| Phoridae | Family | Diptera |
| Phryganeidae | Family | Trichoptera |
| Platygastridae | Family | Hymenoptera |
| Polycentropodidae | Family | Trichoptera |

| Taxa | Taxonomic level | Order |
|---|---|---|
| Pompilidae | Family | Hymenoptera |
| Psocoptera | Order | |
| Psychodidae | Family | Diptera |
| Pteromalidae | Family | Hymenoptera |
| Rhaphidophoridae | Family | Orthoptera |
| Saldidae | Family | Hemiptera |
| Scatopsidae | Family | Diptera |
| Scelionidae | Family | Hymenoptera |
| Sciaridae | Family | Diptera |
| Sciomyzidae | Family | Diptera |
| Scirtidae | Family | Coleoptera |
| Scraptiidae | Family | Coleoptera |
| Sepsidae | Family | Diptera |
| Sisyridae | Family | Neuroptera |
| Sphecidae | Family | Hymenoptera |
| Staphylinidae | Family | Coleoptera |
| Stratiomyidae | Family | Diptera |
| Syrphidae | Family | Diptera |
| Tabanidae | Family | Diptera |
| Tachinidae | Family | Diptera |
| Throscidae | Family | Coleoptera |
| Thysanoptera | Order | |
| Tipulidae | Family | Diptera |
| Trichogrammatidae | Family | Hymenoptera |
| Ulidiinae | Subfamily | Diptera |

Appendix D. Aquatic invertebrate taxa identified in the ¼ $m^2$ vegetation quadrats collected from herbicide-treated, *P. australis*, and remnant marsh sites from Long Point, ON in May 2018.

| Taxa | Taxonomic level | Order |
|---|---|---|
| Arachnida | Subclass | |
| Amphipoda | Order | |
| Araneae | Order | |
| Bivalvia | Class | |
| Caenidae | Family | Ephemeroptera |
| Cecidomyiidae | Family | Diptera |
| Ceratopogonidae | Family | Diptera |
| Chironomidae | Family | Diptera |
| Coenagrionidae | Family | Odonata |
| Collembola | Order | |
| Corduliidae | Family | Odonata |
| Crambidae | Family | Lepidoptera |
| Culicidae | Family | Diptera |
| Curculionidae | Family | Coleoptera |
| Brachycera | Suborder | |
| Dolichopodidae | Family | Diptera |
| Dytiscidae | Family | Coleoptera |
| Ephydridae | Family | Diptera |
| Gastropoda | Class | |
| Lumbriculata | Subclass | |
| Hydrophilidae | Family | Coleoptera |
| Hydrozoa | Class | |
| Isopoda | Order | |
| Lampyridae | Family | Coleoptera |
| Leptoceridae | Family | Trichoptera |
| Limnephilidae | Family | Trichoptera |
| Nematoda | Phylum | |
| Oligochaeta | Subclass | |
| Ostracoda | Class | |
| Platyhelminthes | Phylum | |
| Pleidae | Family | Hemiptera |
| Sciomyzidae | Family | Diptera |
| Scirtidae | Family | Coleoptera |
| Staphylinidae | Family | Coleoptera |
| Stratiomyidae | Family | Diptera |
| Thysanoptera | Order | |
| Tipulidae | Family | Diptera |

Appendix E. Taxa included in Procrustes test analysis from the emergence trap samples (emerging invertebrate samples) and the aquatic invertebrate samples (aquatic invertebrate samples). Taxa were identified to lowest feasible taxonomic level.

| Taxa | Samples |
|---|---|
| Caenidae | Emergence trap |
| Cecidomyiidae | Emergence trap |
| Ceratopogonidae | Emergence trap |
| Chironomidae | Emergence trap |
| Coenagrionidae | Emergence trap |
| Crambidae | Emergence trap |
| Culicidae | Emergence trap |
| Dolichopodidae | Emergence trap |
| Ephydridae | Emergence trap |
| Lampyridae | Emergence trap |
| Leptoceridae | Emergence trap |
| Limnephilidae | Emergence trap |
| Sciomyzidae | Emergence trap |
| Stratiomyidae | Emergence trap |
| Tabanidae | Emergence trap |
| Tipulidae | Emergence trap |
| Dixidae | Emergence trap |
| Hydroptilidae | Emergence trap |
| Lestidae | Emergence trap |
| Libellulidae | Emergence trap |
| Phryganeidae | Emergence trap |
| Polycentropodidae | Emergence trap |
| Chaoboridae | Emergence trap |
| Syrphidae | Emergence trap |
| Ichneumonidae | Emergence trap |
| Scelionidae | Emergence trap |
| Mymaridae | Emergence trap |
| Eulophidae | Emergence trap |
| Pteromalidae | Emergence trap |
| Phoridae | Emergence trap |
| Cosmopterigidae | Emergence trap |
| Caenidae | Aquatic sample |
| Cecidomyiidae | Aquatic sample |
| Ceratopogonidae | Aquatic sample |
| Chironomidae | Aquatic sample |
| Coenagrionidae | Aquatic sample |

| Taxa | Samples |
|---|---|
| Crambidae | Aquatic sample |
| Culicidae | Aquatic sample |
| Dolichopodidae | Aquatic sample |
| Ephydridae | Aquatic sample |
| Lampyridae | Aquatic sample |
| Leptoceridae | Aquatic sample |
| Limnephilidae | Aquatic sample |
| Sciomyzidae | Aquatic sample |
| Stratiomyidae | Aquatic sample |
| Tipulidae | Aquatic sample |
| Tabanidae | Aquatic sample |
| Corduliidae | Aquatic sample |

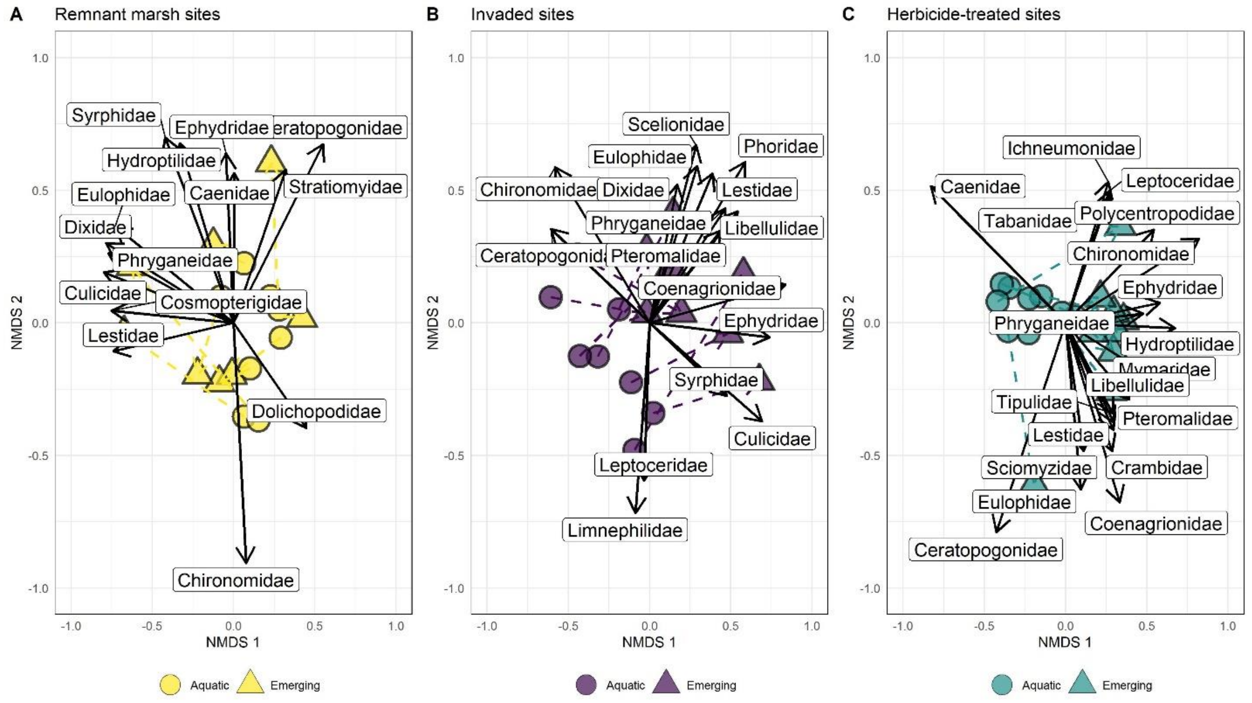


Appendix F. NMDS ordination of emerging and aquatic invertebrate communities present in each habitat. The uninvaded NMDS had a 2D final solution with a stress of 0.161 after 88 iterations and a non-metric $r^2$ of 0.974 (A), the invaded emerging invertebrate NMDS had a 2D final solution with a stress of 0.138 after 48 iterations and a non-metric $r^2$ of 0.981 (B), and the herbicide-treated sites had a 2D final solution with a stress of 0.080 after 49 iterations and a non-metric $r^2$ of 0.993 (C). Aquatic invertebrates were collected in mid-May 2018 from submersed aquatic vegetation and emerging invertebrates were collected from 05-June-18 to 23-July-18. Reasonably correlated taxa ($r^2 > 0.30$) are included as vectors and ellipses are 90% CI.
